# Severe malarial anaemia is associated with parasite enrichment and host alterations in the bone marrow

**DOI:** 10.64898/2026.04.27.720990

**Authors:** Joao da Silva Filho, Elamaran Meibalan, Himanshu Gupta, Dario Beraldi, Selina Bopp, Dyann F. Wirth, Christopher A. Moxon, Eusebio Machete, Cinta Moraleda, Ruth Aguilar, Alfredo Mayor, Dan Milner, Clara Menendez, Matthias Marti

## Abstract

**Background:** Malaria poses a significant global health challenge, with over 200 million infections and around 500,000 deaths annually, predominantly affecting children under 5 in sub-Saharan Africa. Severe malarial anaemia (SMA), a major contributor to morbidity and mortality in this demographic, results from various factors including red blood cell destruction and immune-mediated clearance. The role of these mechanisms in SMA differs based on age and infection history. Recent studies indicate increased accumulation of *P. falciparum* in the bone marrow and spleen, highlighting the need for understanding localized host-parasite interactions.

**Methods:** This study examines the bone marrow response to malarial infection in young children with SMA or mild malarial malaria in Mozambique, contrasting it with responses observed in peripheral blood in the two cohorts.

**Results and conclusions:** The study demonstrated that SMA is associated with increased red blood cell production, iron metabolism and tissue injury, as well as higher total parasite biomass including peripheral and bone marrow parasitaemia. We also demonstrate that direct analysis of bone marrow aspirates provides far more resolution in stratifying host signatures and drivers of malarial anaemia across a spectrum of severity than systemic measures from blood samples.

## Background

Malaria remains a major global health issue, with over 200 million new infections and about half million fatal cases each year. Most of the fatal cases are in sub-Saharan Africa, in particular in children below the age of 5 (WHO, 2024). Major clinical syndromes caused by *P. falciparum* infection include cerebral malaria, respiratory distress and severe malarial anaemia (SMA).

In young children SMA is a major cause of morbidity and mortality. SMA has a complex aetiology, including red blood cell (RBC) destruction due to parasite lysis and splenic clearance (also of uninfected RBCs), immune-mediated clearance (through complement, antibodies, neutrophils or macrophages), as well as depleted iron stores and dyserythropoiesis in the bone marrow. The contribution of these processes to SMA varies depending on the children’s age and exposure to infection. In children with repeated or chronic infections anaemia is largely driven by bone marrow dyserythropoiesis and iron depletion, while splenic and immune-mediated clearance plays a major role in acute infection with high parasite load (Buffet *et al*, 2010; Moxon *et al*, 2020; White, 2018). Most of these observations were made in a few cohorts from sub-Saharan Africa, and it remains to be shown whether this aetiology also applies more broadly to sub-Saharan Africa and to other malaria endemic regions in the world.

Recent studies have demonstrated major accumulation of *P. falciparum* blood stages in the bone marrow and spleen (Aguilar *et al*, 2014a; De Niz *et al*, 2018; Joice *et al*, 2014; Kho *et al*, 2021a; Kho *et al*, 2021b), suggesting specific interactions and physiological effects in these organs that may contribute to SMA. We hypothesised that the severity of malarial anaemia in young children is driven primarily by bone marrow-specific host-parasite interactions rather than systemic responses detectable in peripheral blood. To test this hypothesis, we specifically investigated the host response to bone marrow infection in comparison to peripheral blood in a cohort of young children with SMA in Mozambique.

## Methods

### Ethics statement

The original study was carried out at the Centro de Investigação em Saúde de Manhiça in the Manhiça District of southern Mozambique (Aguilar *et al*., 2014a; Aguilar *et al*, 2014b). The study protocol was approved by the National Committee for Bioethics in Health of Mozambique, CNBS (259/CNBS) and the Hospital Clínic of Barcelona Ethics Review Committee (2008/4089). Individual written informed consent was obtained from the parents/guardians of all children before enrolment. All children received treatment according to national guidelines.

### Clinical parameters and sample formats

From the cohort as described previously (Aguilar *et al*., 2014a; Aguilar *et al*., 2014b), a total of 31 patients were included (see **Table S1** for patient meta data). For all patients, we had matched peripheral blood and bone marrow samples. Details of peripheral blood and bone marrow aspirate sample collection are described in the original studies (Aguilar *et al*., 2014a; Aguilar *et al*., 2014b). We used blood plasma for Luminex arrays and analysis of some individual biomarkers, and cellular pellets from blood (PBMCs) and bone marrow aspirates (BMMCs) for qRT-PCR and Nanostring arrays (**Table S1**).

### Analysis of protein biomarkers from blood plasma

Blood plasma from peripheral blood collected and stored as described (Aguilar *et al*., 2014a; Aguilar *et al*., 2014b) was used to measure levels of prealbumin (detail) and lactate dehydrogenase (LDH) with a Vitros DT 6011 Chemistry analyzer (Siemens Healthcare). In addition, C-reactive protein (CRP) and erythropoietin (EPO) levels were already analysed from blood plasma (Aguilar *et al*., 2014a). We also measured parasite LDH levels as a proxy for parasite total parasite biomass (Barber *et al*, 2015). To measure pLDH in peripheral blood plasma samples, ELISA was performed using a matching pair of capture and detection antibodies. Briefly, a 96-well microtiter plate was coated with mouse monoclonal anti-*Plasmodium* LDH (MyBioSource, clone #M77288; MBS838698 at a concentration of 4 μg/ml in PBS (pH 7.4) and incubated overnight at 4°C. The plate was washed and incubated with blocking buffer (PBS-BSA 1% - reagent diluent) at RT for 2 h. After washing, samples were diluted 1:4, added to the plate, and incubated for 2 h. Next, plates were washed, and HRP-conjugated anti-pLDH detection antibody (clone #M12299, MBS833805), diluted 1:1,000 in blocking buffer, was incubated for 1h at RT. Plates were washed and incubated for 15 min with substrate solution (OPD); the reaction was stopped by adding 2.5 M sulfuric acid. Optical density was determined at 450 nm. The cutoff of positivity was defined by correcting absorbance values generated in the plasma samples from blank values (plate controls). The total protein concentration from *P. falciparum* schizont extracts was determined, and samples were used to perform standard curves ranging from 15,625 ng/ml to 2,000 ng/ml. Lower absorbance values were in the range of O.D = 0.004–0.007. All positive samples gave O.D. values equal to or higher than 0.15.

The concentrations of selected proteins in the plasma (peripheral blood protein markers) were quantified using a customized multiplex bead (Luminex) assay (R&D Systems) containing the following 36 analytes (**Table S2**): EC activation and damage: Angiopoietin-1 (Ang-1), Angiopoietin-2 (Ang-2), E-Selectin/CD62E, ICAM1, VCAM1, Syndecan-1, VEGF; Platelet activation and procoagulation markers: P-selectin/CD62P, IL-11, Thrombopoietin (TPO), PAI-1; Chemokines: CCL2/MCP-1, CCL5/RANTES, CCL11/Eotaxin, CXCL12-SDF1; Myeloid response: Granulocyte colony-stimulating factor (G-CSF), Granulocyte-macrophage colony-stimulating factor (GM-CSF), Macrophage colony-stimulating factor (M-CSF), IL-27; Cytokines: TNF-α, IL-3, IL-6, IL-7, IL-8/CXCL8, IL-10, IL-15, IL-21; IL-1 family and Inflammasome response: IL-1α, IL-1β; Type I and II Interferons (IFNs): IFN-α, IFN-β, IFN-γ; Hematopoiesis-related cytokines: Osteopontin (OPN), Stem cell factor (SCF/c-kit Ligand), c-Kit, Flt-3l.The assays were conducted according to the manufacturer’s instructions in a Bio-Rad Bio-Plex 200 Systems. Fluorescence intensity data from the assays were used for further analysis.

### Analysis of transcript biomarkers by qRT-PCR and/or Nanostring nCounter array

Aliquots of RNA samples from PBMCs or bone marrow aspirates collected and stored as described (Aguilar *et al*., 2014a) were used for analysis of transcript biomarkers. cDNA was synthesized with the High-Capacity cDNA Reverse Transcription Kit with RNase Inhibitor (Thermofisher - cat number: 4374966), following the manufacturer protocol, and mRNA expression of genes were determined by qRT-PCR. For qRT-PCR, we selected a subset of the protein biomarkers analysed from blood plasma that were not present in the host nCounter panel (**Table S2**). Primers were purchased as TaqMan Gene Expression Assay (FAM) from ThermoFisher (4331182): EPO (assay ID: Hs01071097_m1); CCL5/RANTES (assay ID: Hs00982282_m1); CCL2/MCP-1 (assay ID: Hs00234140_m1); IFNG (assay ID: Hs00989291_m1); IFNB1 (assay ID: Hs01077958_s1); CSF3 (assay ID: Hs00738432_g1); VEGFA (assay ID: Hs00900055_m1); SELP (assay ID: Hs00927900_m1); SDC1 (assay ID: Hs00896424_g1); VCAM1 (assay ID: Hs01003372_m1); ANGPT2 (assay ID: Hs01048042_m1). Human GAPD (GAPDH) Endogenous Control (FAM™/MGB probe, non-primer limited, 4333764T) was used as housekeeping gene. Real-time qRT-PCR was performed on an ViiA 7 Real-Time PCR System (Applied Biosystems). Cycling parameters were 95 °C for 20 s and then 40 cycles of 95 °C (1 s) and 60 °C (20 s), followed by a melting curve analysis. The median cycle threshold (Ct) value and 2 -ΔΔCt method were used for relative quantification analysis, and all Ct values were normalized to the GAPDH mRNA expression level using the qPCRtools R package (10.3389/fgene.2022.1002704). Results expressed as means and SEM of biological replicates are shown. In addition to qRT-PCR, we also employed the probe-based Nanostring nCounter array platform (Bruker Technologies). The probe set targets a total of 37 transcript biomarkers and 10 housekeeping genes (**Table S2**). The probe set also targets a total of 456 parasite genes (**Table S2**) that have been validated in a series of studies analysing blood and tissue samples from malaria patients (De Niz *et al*., 2018; Meibalan *et al*, 2021; Van Tyne *et al*, 2014). Nanostring raw counts were first normalised using the NanoStringNorm function (R package NanoStringNorm 1.2.1) with options: CodeCount = ‘sum’, Background = ‘mean’, SampleContent = ‘housekeeping.sum’. We further normalised the dataset using the function normalizeQuantileRank.matrix (R package aroma.light 3.16) so that all markers have the same distribution across samples.

### Data analysis

#### Statistical analysis

To ensure that the differences observed between patient samples were due to severity of malarial anaemia and not due to differences in age, self-reported sex or comorbidities (fixed effects), their effect was tested using linear models according to the function lmer of the lme4 R package version 1.1.32) (Douglas Bates, 2015), with the following formula: lm[variable ∼ age + gender + comorbidities, data = dataset]. For parameters with comorbidity (i.e., acute epstein-barr virus infection, acute parvovirus B19 infection, albumin deficiency, malnutrition, dyserythropoiesis, thalassemias), sex, or age influence, estimates of predicted influence (fitted values) were subtracted from the raw parameter values, and residuals were used for downstream statistical testing. Data normality was checked by the results of the D’Agostino and Pearson, Anderson-Darling, Kolmogorov-Smirnov, and ShapiroWilk tests. Student’s t test was used to compare medians between groups (severe *versus* moderate malarial anaemia) with normally distributed data, and datasets with non-normal distributions were compared using the Mann-Whitney test. Fisher’s exact test was used for categorical data. All tests were performed two-sided using a nominal significance threshold of *p* < 0.05, unless otherwise specified. When appropriate to adjust for multiple hypothesis testing, Bonferroni, FDR, or Benjamini-Hochberg (BH), two-stage step-up (https://www.mathworks.com/matlabcentral/fileexchange/27423-two-stage-benjamini-krieger-yekutieli-fdr-procedure) correction methods were applied to test significance at *p* <0.05 threshold. Analyses were performed and the graphs were generated in GraphPad Prism 9 (version 9.5.1 [528], 2023) and RStudio software (version 2023.03.0+386; 2023).

#### Dimensionality reduction by Principal Component Analysis (PCA) and Exploratory Factor Analysis (EFA)

We carried out PCA on the normalized data using the PCA function of the FactoMineR R package (v 2.8) with default parameters (Husson *et al*, 2017) For each principal component (PC), we determined the most significantly *(p* < 0.05) associated variables with a given principal component (using the function dimdesc), and the contribution of each variable for each PC (using the function get_pca_var). For visualization of PCA results, ggplot2 (v 3.4.2), factoextra (v 1.0.7), and corrplot (v 0.92) R packages were used. EFA was used to reduce the dimensionality of the clinical and bone marrow markers (the latter derived from the nCounter and qRT-PCR experiments) into a smaller set of features, without losing information, to identify the driving “signatures” of variance according to severity of malarial anaemia following the same analysis pipeline previously published (Da Silva Filho *et al*, 2024). The factor loading values of each clinical parameter and bone marrow marker was used to determine closest association with a particular factor (which we called clinical and bone marrow (BM) signatures). For instance, clinical parameters with no association with a corresponding signature are expected to have factor loadings close to zero, whereas parameters with strong association are expected to have large absolute loading values. The composition of each clinical and BM signature - after sorting parameters across signatures based on the factor loadings – was then plotted in heatmaps, and the mean ± SEM of the sample loading scores per patient, grouped according to disease group, was represented in box plots. The sign of the sample loading score indicates the direction of the effect: samples with high positive score values indicate that the parameters composing that clinical or bone marrow signature are enriched in those samples, whereas samples with high negative values indicate reduced values for the parameters composing that clinical or peripheral blood signature.

Ensembl (http://www.ensembl.org), Metascape (http://metascape.org) and PantherDB (https://geneontology.org) pathway analyses were performed to extract the biological processes associated with each marker in each bone marrow signature. These methods support gene identifier types, such as Entrez Gene ID, Ensembl ID, RefSeq and UniProt ID. Gene lists (encoding specific cytokines) were tested by custom analysis. Each method outputs a results page, displaying the lists of shared GO terms used to describe the specific set of genes, an indication of over/underrepresentation for each term, and a *p* value. The GO result lists from each database, for each set of cytokines, according to the peripheral blood signature they belong to, is included in the **Tables S3** and **S4**.

#### Multi-modal data integration

Our dataset includes parallel characterization of clinical, host and parasite signatures in the PB and BM, i.e., clinical parameters, BM and PB host protein and RNA profiling, stage quantification from bone marrow and blood smears and total parasite biomass from pLDH ELISA. This rich dataset offers the opportunity to analyse signatures (clinical and molecular) and their tissue source (BM *vs* PB) to define the responses driving the severity in malarial anaemia. We integrated the clinical data, parasite biomass data, as well as PB and BM host protein and RNA profiling using unsupervised integrative approaches to identify latent factors of variation in multi-modal high-dimensional data (Argelaguet *et al*, 2020; Argelaguet *et al*, 2018; Chang *et al*, 2021; Fanaee & Thoresen, 2019; Hore *et al*, 2016; Josse & Husson, 2016; Pagès, 2014; Velten *et al*, 2022). This integration aims to disentangle the host and parasite driving sources of variation between the SMA and MMA. We first applied Multiple Factor Analysis (MFA) to integrate clinical data, peripheral blood biomarker profiling and bone marrow features using the function MFA in the missMDA v 1.18 R package)(Pagès, 2014), following a previously described pipeline (Da Silva Filho *et al*., 2024). In brief, MFA consists of performing a global PCA on the dataset concatenating the weighted matrix of each source of variables. Here, we applied MFA with the following aims: (i) to investigate the differences and similarities between patients from a multidimensional point of view, as well as the correlation between variables; (ii) to highlight similarities and differences between groups of variables (pointing out what is common *versus* specific between MMA and SMA); (iii) finally, MFA was also used to balance the influence of the groups of variables (clinical, peripheral blood biomarkers and bone marrow features) in the analysis in such a way that no single group, with correlated variables for instance, dominates the top dimensions of variability. For this purpose, the MFA approach calculates for each group of variables (clinical, peripheral blood and bone marrow-derived) a principal component (PC) and then each PC value is divided by the square root of the first eigenvalue. To indicate the source of the assay of each variable, they were grouped as clinical, parasite, peripheral blood biomarker and bone marrow biomarker. The data for each source was centred and scaled, and missing values were imputed using the function imputeMFA (missMDA R package).

Next, we used Multi-Omics Factor Analysis (MOFA) (Argelaguet *et al*., 2020; Fanaee & Thoresen, 2019; Josse & Husson, 2016) another computational method to integrate the multiple data modalities, including the parasite stage distribution derived from transcript profiling, in an unsupervised fashion to disentangle the principal drivers of variation (here called “integrated signatures”) between MMA and SMA, as previously described (Da Silva Filho *et al*., 2024).The method was designed for integrating data modalities by a common sample space (measurements derived from the same set of samples), where the features (variables) are distinct across data modalities (approaches). Given several data matrices with measurements of parameters from different approaches on the same or on partially overlapping sets of samples, MOFA infers an interpretable low-dimensional data representation in terms of (hidden) factors here to referred as “integrated signatures”. MOFA then disentangles to what extent each factor is either unique to a single data modality or representative of variation from multiple modalities, thereby revealing shared axes of variation between the different layers. After model training, each factor (i.e., integrated signature) captures a different source of variability in the data, as they are defined by a linear combination of the input features. A good sanity check for model quality control is to verify whether the factors are largely uncorrelated which confirms a good model fit and suggests that the chosen number of factors in the model is appropriate and the normalization is adequate. The variance decomposition by factor (or integrated signature) was also evaluated, as it summarizes the percentage of variance explained by each factor across each data modality (clinical, parasite transcriptional staging, peripheral blood protein marker, and bone marrow marker) from the heterogeneous dataset.

## Results

### Stratification of patient cohort into moderate and severe malarial anaemia based on differences in clinical parameters

We followed a retrospective cohort of hospitalized paediatric patients admitted at the Centro de Investigação em Saúde de Manhiça in the Manhiça District of southern Mozambique. Children (n=31) with confirmed *Plasmodium falciparum* infection, aged between 1-59 months with haemoglobin levels of <11g per dL and bone marrow aspiration within the past four weeks were recruited between 2008 and 2010 (**Table S1**). Original studies describing this cohort focused on comparative analysis of blood and bone marrow enrichment of asexual and sexual *P. falciparum* stages (Aguilar *et al*., 2014a), and the association of *P. falciparum*-derived hemozoin in the bone marrow with severity of anaemia (Aguilar *et al*., 2014b). For our analysis, we divided patients into two groups of malarial anaemia severity (severe malarial anaemia – SMA, n=17, and moderate malarial anaemia – MMA, n=14), according to the haemoglobin concentration cut-off at 7g/dL (WHO Team: Monitoring and Surveillance Nutrition and Food Safety (MNF), 2011). All clinical, parasitological and molecular parameters have been adjusted by age and gender using linear regression models. Initial comparisons of clinical parameters according to this patient stratification showed significant increase in i) erythropoietin (EPO) plasma levels indicating stimulating of red blood cell production, ii) tissue injury markers (e.g., human lactate dehydrogenase (LDH) and C-reactive protein, CRP), iii) total parasite biomass (represented by parasite LDH concentration, pLDH (Barber *et al*., 2015)), as well as iv) peripheral blood (PB) and bone marrow (BM) parasitaemia in SMA compared to MMA patients (**Figure 1A**). To disentangle the cohort heterogeneity further and to characterize the drivers of these 2 clinical trajectories in malarial anaemia, we performed principal component analysis (PCA) and Exploratory Factor Analysis (EFA) with the clinical parameters determined at hospitalisation. With these unsupervised approaches, we were able to confirm the supervised grouping of malaria patients in those with severe *versus* those with moderate anaemia (**Figures 1B, C**). Furthermore, both PCA and EFA validated the increase in EPO plasma levels, iron metabolism (i.e., ferritin and plasma iron), increase in tissue injury markers (host LDH and CRP), and increased total parasite biomass (i.e., pLDH) in SMA patients compared to moderate malarial anaemia cases (represented by clinical signature 1) (**Figure 1C,D**). Interestingly, both PCA and EFA also indicated increased bone marrow gametocytaemia in SMA patients (**Figure 1C**). Altogether parasite and host factors, in particular responses developing in the bone marrow, seem to be contributing to the severity of malarial anaemia.

**Figure 1.**
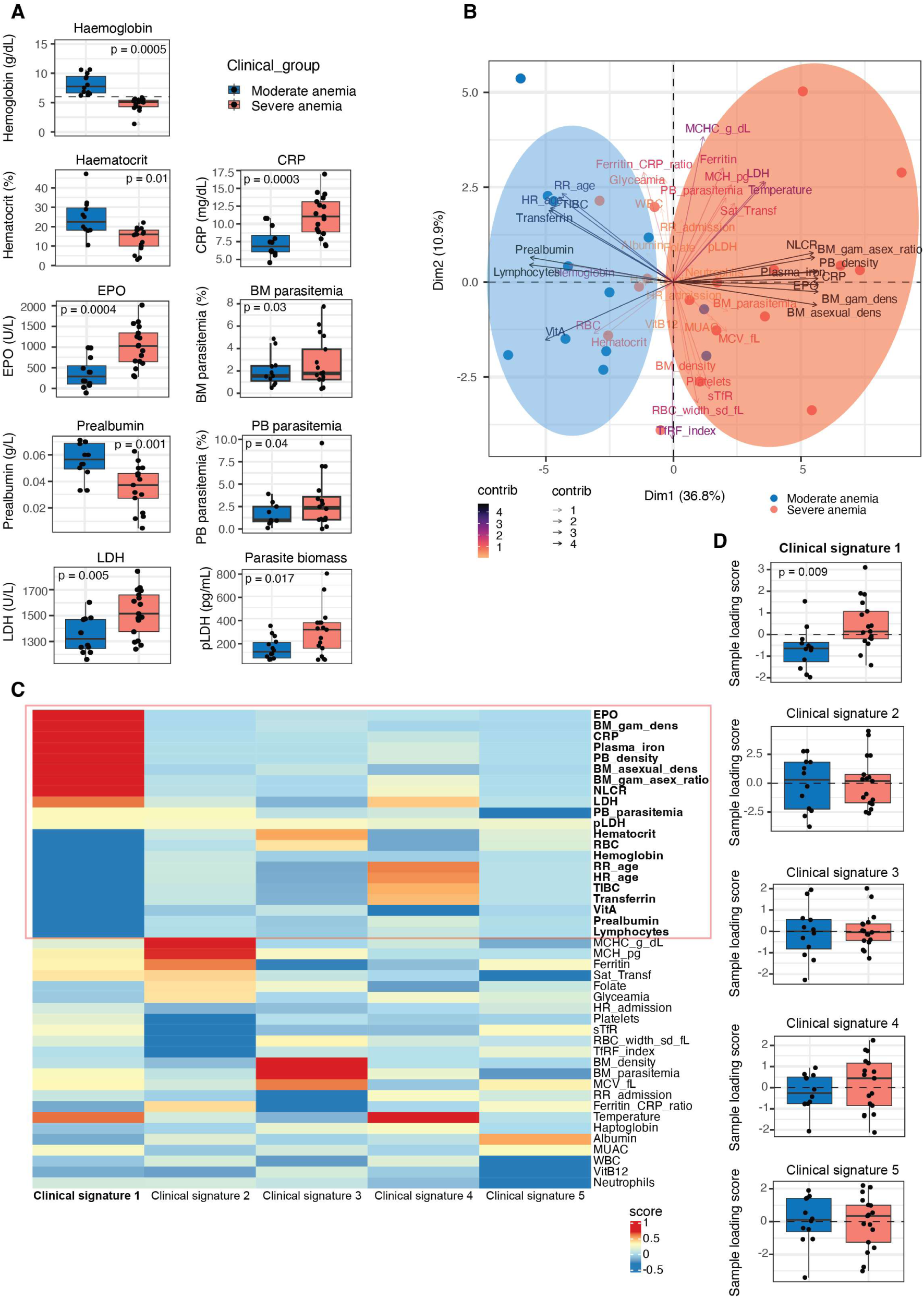
Stratification of patient cohort in MMA and SMA based on differences in clinical parameters. **A.** Stratification of patient cohort into two clinical groups - moderate (MMA) *versus* severe anaemia (SMA). Shown are those parameters with a significant difference between SMA and AMA (*p*<0.05). **B,C.** Principal component analysis (PCA) (**B**) and Exploratory Factor Analyis EFA (**C**). EFA groups clinical parameters into five factors termed clinical signatures 1-5. Clinical signature 1 (red box) is significantly different (*p*<0.05) between SMA and SMA (**C**), confirming separation by PCA (**B**, red *versus* blue circles) and of individual parameters in **A**.

### BM but not PB host biomarkers are associated with SMA

To further investigate whether host signatures drive the severity in malarial anaemia in our cohort, we performed multiplex profiling using a bead-based luminex assay to quantify 36 host protein biomarkers (**Table S2 and Figure S1**) and transcript profiling (**Figure 2A,B** and **S2**) of samples collected at hospital admission. No bone marrow supernatants were available for luminex, hence we assessed a subset of these markers and an additional panel on a transcript level by qRT-PCR (representing 11 luminex markers) and a probe-based platform (nCounter, representing 6 luminex markers and 41 additional host markers) using bone marrow mononuclear cells (BMMCs). Out of 36 protein markers measured by Luminex in PB, only Syndecan-1 (a marker of endothelial cell damage), TNF-a and IL-27 (markers related to myeloid activation) were significantly increased in SMA compared to MMA patients (**Figure S1**). Similarly, qRT-PCR analysis in peripheral blood mononuclear cells (PBMCs) did not show major associations with severity of malarial anaemia (**Figure S2A**). Indeed, PCA performed with the PB Luminex (**Figure S1**) and PB qRT-PCR data (**Figure 2A**) did not cluster the patients according to severity of malarial anaemia.

**Figure 2.**
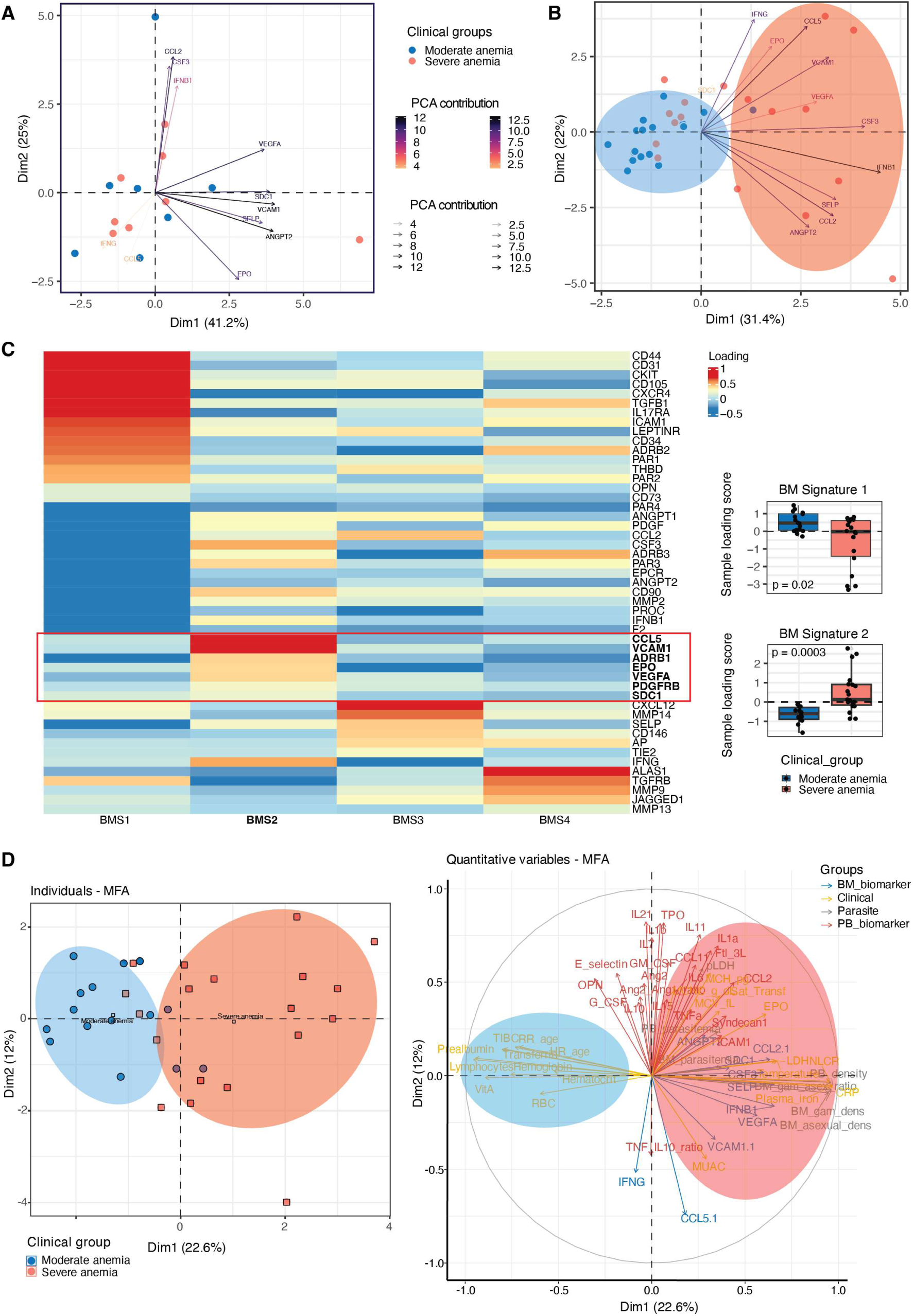
BM but not PB host biomarkers are associated with SMA. Stratification of qRT-PCR immune marker panels in PB samples (**A**) and BM samples (**B**) between SMA and MMA patients**. C.** EFA analysis of qRT-PCR and nCounter transcript quantification in BMMCs. EFA groups transcript markers into four factors termed bone marrow signatures 1-4. BM signatures 1 and 2 (red box) are significantly different (*p*<0.05) between SMA and MMA patients. **D.** Multiple Factor Analysis (MFA) to integrate the clinical data, PB and BM profiling. PCA of the MFA output stratified the patients based on the MMA vs SMA grouping.

In contrast, quantification of the qRT-PCR markers in BMMCs showed upregulation of *SDC1* (encoding Syndecan-1), *VCAM1, VGFA* (encoding markers related to endothelial cell activation and damage), *CCL5* (an immune cell chemokine), *CSF3* (encoding G-CSF protein, a marker related to myelopoiesis) and *IFNB1* (indicative of IFN-mediated responses) in the SMA patients compared to the MMA patients (**Figure S2B**). PCA based on nCounter transcript quantification in BMMCs revealed a clear separation of SMA and MMA cases (**Figure 2B**). In addition, EFA of BM signatures (based on combined qRT-PCR and nCounter transcript quantification in BMMCs) corroborated the increase of genes related to erythropoiesis, endothelial cell activation and damage, immune chemoattraction and myeloid response in the bone marrow of SMA patients, as shown for the BM signature 2 (**Figure 2C, Tables S3 and S4**).

Our dataset includes parallel characterization of clinical, host and parasite signatures in the PB and BM, i.e., clinical records, BM and PB host protein and RNA quantification, stage quantification from bone marrow and blood smears and total parasite biomass from pLDH ELISA. This offers the opportunity to analyse signatures (clinical and molecular) and their tissue source (BM *vs* PB) to define the responses driving the severity in malarial anaemia. We integrated the clinical data, parasite biomass-related data, as well as PB and BM host protein and transcriptomic profiling using unsupervised integrative approaches to identify latent factors of variation in multi-modal high-dimensional data (Argelaguet *et al*., 2020; Argelaguet *et al*., 2018; Chang *et al*., 2021; Fanaee & Thoresen, 2019; Hore *et al*., 2016; Josse & Husson, 2016; Pagès, 2014; Velten *et al*., 2022). This integration disentangles the host and parasite driving sources of variation between the SMA and MMA. Specifically, we applied Multiple Factor Analysis (MFA), which clearly stratified patients based on malarial anaemia severity **(Figure 2D, left panel)**. Clinical parameters, such as LDH, plasma iron, transferrin, CRP, EPO, haemoglobin, haematocrit, bone marrow host signatures, such as *IFNB1, CSF3*, and *SDC1* transcript levels, and total parasite biomass (sexual and asexual densities) in the BM were the main host and parasite factors driving the stratification between the 2 groups according to malarial anaemia severity (**Figure 2D right panel, Figure S3**).

### Immature gametocyte signatures are increased in BM compared to PB

In parallel to the host protein and transcript profiling, we also investigated the parasite transcriptional profiles driving the severity in malarial anaemia in our cohort. For this purpose, we analysed *P. falciparum* gene expression from BM and PB using an nCounter probe array targeting a total of 456 parasite genes (De Niz *et al*., 2018; Meibalan *et al*., 2021; Van Tyne *et al*., 2014). PCA based on parasite gene expression in red blood cells from BM and PB did not stratify patient samples according to malarial anaemia severity (**Figure 3A**), suggesting that parasite expression patterns in the BM (and blood) of SMA patients are not significantly different from the parasite expression patterns in the BM of MMA patients. Indeed, differential parasite gene expression analysis between matched BM and PB samples and between SMA and MMA, respectively, revealed minimal variation (**Figure 3B, Table S5**).

**Figure 3.**
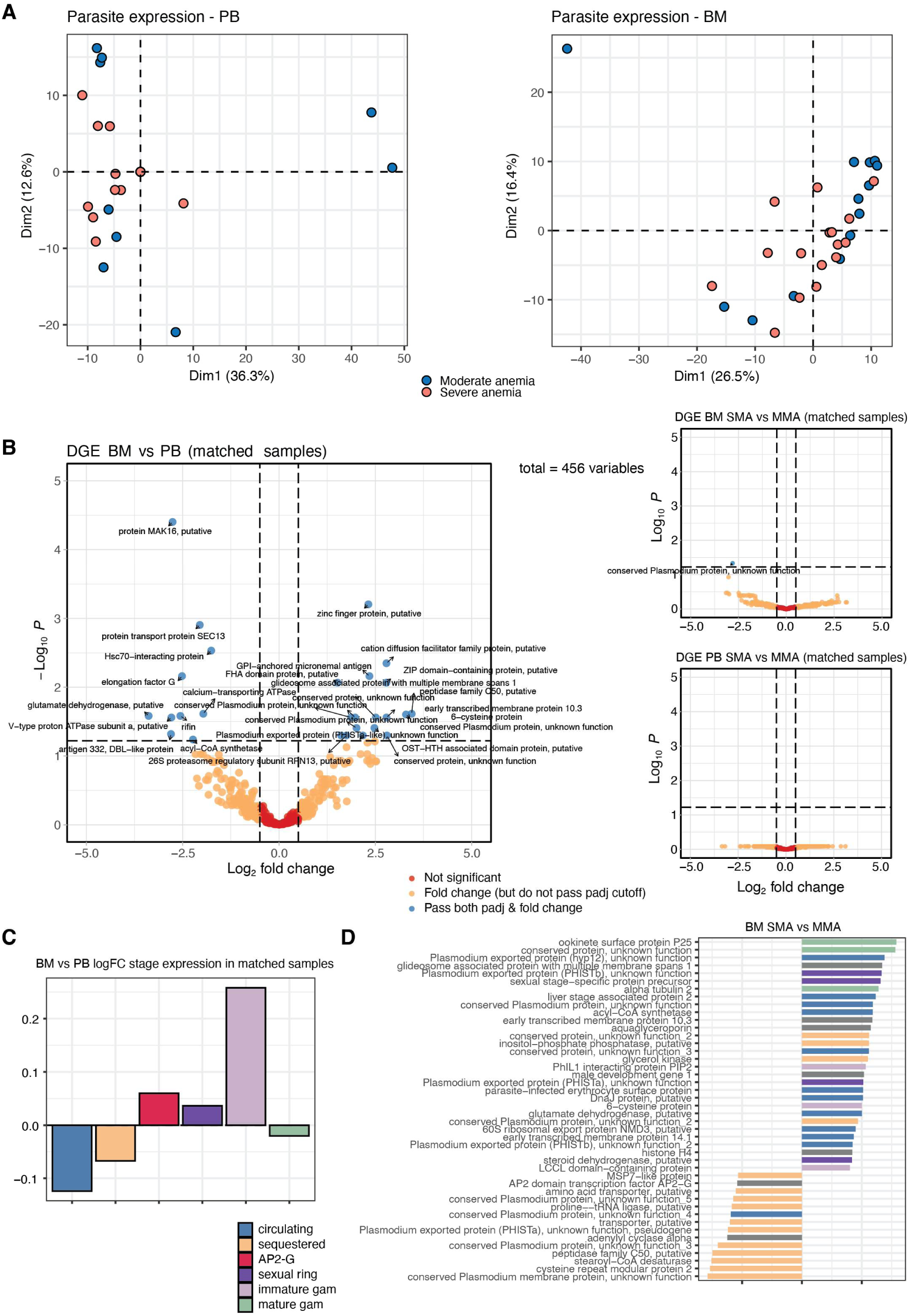
Immature gametocyte levels are increased in BM. **A.** PCA based on parasite gene expression from nCounter transcript signatures in PBMCs (left) and BBMCs (right). **B.** Volcano plots showing differential parasite gene expression between BM and PB from all patient-matched samples (left) and stratified by SMA *versus* MMA (right). **C.** Difference in parasite stage composition between BM and PB in patient matches samples determined using gene annotations per stage from (Pelle *et al*., 2015). **D.** Difference in BM gene expression between SMA and MMA patients.

However, once gene expression was stratified by parasite stage (as previously defined (Pelle *et al*, 2015)), major differences in the patterns of parasite transcriptional signatures could be observed when comparing matched BM and PB samples (**Figure 3C**). Specifically, this analysis revealed enrichment of *P. falciparum* markers for forming (*ap2-g*) and developing gametocytes (sexual ring and immature gametocytes) in the BM, and enrichment of asexual parasites (circulating and sequestered) and mature gametocytes in the peripheral blood, confirming previous observations that early gametocyte development occurs in the bone marrow (Aguilar *et al*., 2014a; Joice *et al*., 2014). *Ap2-g* expression is also non-significantly elevated in SMA vs MMA patients from BM (**Figure 3D**).

### Multi-modal data integration reveals differences in host biomarkers and parasite stages between MMA and SMA

To integrate clinical parameters, PB and BM host biomarkers (at protein and transcript levels) and also to include parasite signatures from differential parasite staging based on nCounter gene expression (besides stage quantification from bone marrow and blood smears) we applied MOFA (Multi-Omics Factor Analysis) (Argelaguet *et al*., 2020; Argelaguet *et al*., 2018; Velten *et al*., 2022) **(Figure 4)**. In brief, this computational method allows us to further integrate multiple data modalities in an unsupervised fashion to disentangle the principal drivers of variation (here called “integrated signatures 1-3”) between SMA and MMA patients **(Figure 4A, left panel).**

**Figure 4.**
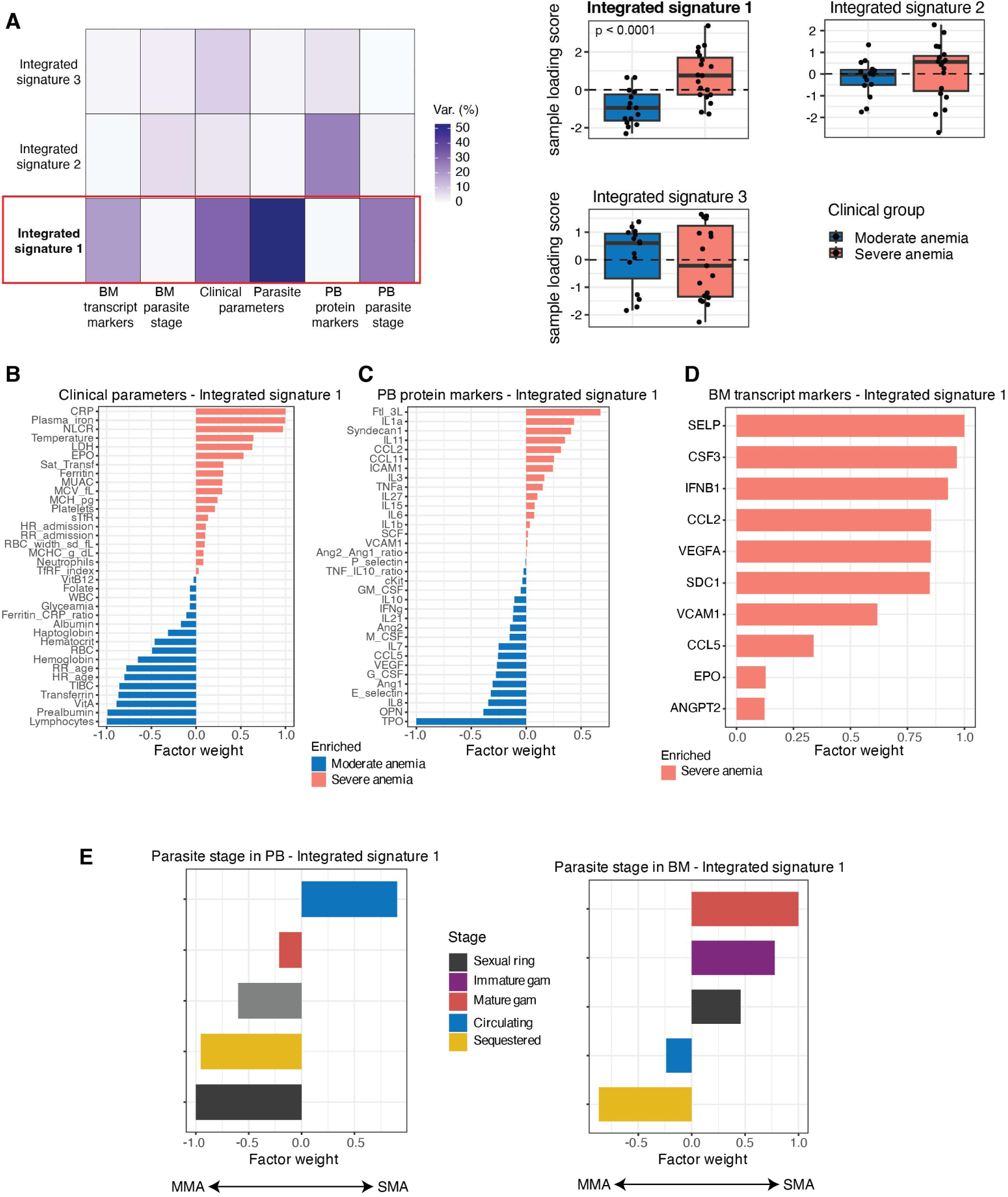
MOFA reveals differences in host biomarkers and parasite stages between MMA and SMA. **A.** MOFA (Multi-Omics Factor Analysis) to integrate clinical parameters, PB and BM host biomarkers and parasite signatures and stratifies them into three integrated signatures (IS1-3). The MOFA factors 1-3 (integrated signatures, IS1-3) capture variation of the total variance across all data modalities per clinical group (SMA *versus* MMA). IS 1 (red box) is significantly different (*p*<0.05) between SMA and MMA patients. **B.** Evaluation of clinical parameters positively or negatively associated with IS1. **C.** Evaluation of PB biomarkers (derived from Luminex) positively or negatively associated with IS1. **D.** Evaluation of BM biomarkers (derived from qRT-PCR data in BMMCs) positively or negatively associated with IS1. **E.** Evaluation of BM and PB parasite parameters (derived from nCounter transcript levels in PBMCs and BMMCs) positively or negatively associated with IS1.

Integrated signature 1 (IS1) was the only factor significantly increased in SMA cases (**Figure 4A, right panel**). In this analysis parameters with positive factor weight values are enriched in SMA cases (thus reduced in MMA cases) and those with negative factor weight values are reduced in SMA cases (thus increased in MMA). Evaluation of clinical parameters (positively or negatively) associated with IS1 confirmed initial EFA based on clinical data, demonstrating enrichment of erythropoiesis (EPO plasma levels), tissue injury (LDH and CRP) and other severe anaemia markers (MCV_fl, plasma iron, sTfR, RBC width, ferritin, etc), as well as reduced RBC, haemoglobin and haematocrit levels in SMA compared to MMA patients (**Figure 4B**). The top PB protein markers (derived from the Luminex data) associated with IS1 (and enriched in the SMA cases) are related to EC activation and damage and leukocyte chemokines (ICAM-1, Syndecan-1, IL-11, CCL2, CCL11) (**Figure 4C**). BM biomarkers (derived from the qRT-PCR data in BMMCs) associated with IS1 (and enriched in SMA compared to MMA cases) include those related to erythropoiesis (EPO), EC activation and damage (VEGFA, VCAM1, SDC1) and immune cell chemoattraction (CCL5, CCL2) (**Figure 4D**). Analysis of parasite stages (derived from nCounter *P. falciparum* transcript expression) associated with IS1, revealed enrichment of gametocyte signatures (mature, immature and sexual rings) in the BM of SMA compared to MMA patients (and correspondingly depletion in PB) (**Figure 4E**).

## Discussion

SMA is the most common complication in *P. falciparum* infection, particularly among children in high transmission settings. Multiple factors contribute to SMA, including haemolysis and immune-mediated clearance of infected and uninfected RBCs, as well as depleted iron stores and bone marrow dyserythropoiesis. Here we aimed to disentangle and quantify the contribution of these factors in the aetiology of SMA. Importantly, we have applied multiplexed protein and transcript analysis from peripheral blood and bone marrow samples and used integrated approaches for a comprehensive analysis of the host and parasite drivers of SMA in a cohort of young children in Moçambique.

Stratifying the patient cohort based on blood haemoglobin levels in patients with severe *versus* moderate anaemia, clinical parameters demonstrated that SMA is associated with increased red blood cell production, iron metabolism and tissue injury, as well as higher total parasite biomass including peripheral and bone marrow parasitaemia. Integrating analysis of host and parasite markers directly from blood plasma and PBMCs and from bone marrow BMMCs enabled us to further untangle the directionality and compartmental resolution of host and parasite parameters in SMA *vs* MMA. Based on analysis of blood plasma and PBMCs only a few differences between SMA and MMA were detected, including increased levels of Syndecan-1 (a marker of endothelial cell damage), as well as TNF-a and IL-27 (markers related to myeloid activation). In contrast, we observed profound differences in BMMC signatures, including increased levels in multiple markers of erythropoiesis, endothelial activation and damage, immune chemoattractants and myeloid response elements, as well as asexual parasite and gametocyte density.

The study has some limitations. First, we only analysed host and parasite signatures in a cohort of SMA and MMA patients as there were no healthy controls included in the original study; hence we are missing the baseline levels for the parameters measured. Second, the lack of cellular or temporal resolution precludes any determination of causality in the observed associations. We therefore propose follow up studies that are performed at single cell level from both compartments and follow patients by repeated blood sampling, as time course sampling of bone marrow material is ethically not justifiable.

## Conclusions

We demonstrate that direct analysis of bone marrow aspirates provides far more resolution in stratifying host signatures and drivers of malarial anaemia across a spectrum of severity than systemic measures from blood samples. Our analysis also confirms parasite enrichment in the bone marrow, and specific increase in gametocyte formation in the bone marrow of SMA compared to MMA patients. As direct bone marrow analysis in clinical settings may not be routinely possible, our data further corroborates previous findings indicating that systemic (i.e. peripheral blood) levels of both host Syndecan-1 and parasite pLDH are predictors of host responses in deep tissues (in particular, the bone marrow) and malaria severity (Barber *et al*, 2021; Barber *et al*., 2015; Silva-Filho *et al*, 2021). Altogether our study links up and confirms various findings relating to malarial anaemia by combining host and parasite signatures with peripheral blood and bone marrow analysis.

## List of abbreviations

SMA: Severe malarial anaemia
MMA: moderate malarial anaemia
RBC: Red blood cell
LDH: lactate dehydrogenase
PBMCs: Peripheral blood mononuclear cells
BMMCs: Marrow mononuclear cells
PCA: Principal Component analysis
MOFA: Multi-OMICs factor analysis

## Declarations

Ethics approvals and consent were obtained and listed under ethics statement. All data generated or analysed during this study are included in this published article and its supplementary information files. The authors declare no conflict of interest.

## Author contributions

JL: Conceptualization, formal analysis, visualization, writing – original draft. EM: Investigation, writing – review and editing. HG: Investigation, writing – review and editing. DB: Formal analysis, visualisation. SB: Investigation. DFW: Supervision. CAM: Conceptualization, writing – review and editing. EM: Investigation. CM: Investigation. RA: Supervision. AM: Conceptualization, supervision, writing – review and editing. DM: Conceptualization, writing – review and editing. CM: Supervision. MM: Conceptualization, supervision, visualization, project administration, writing – review and editing.

## Supplementary figures

**Figure S1.**
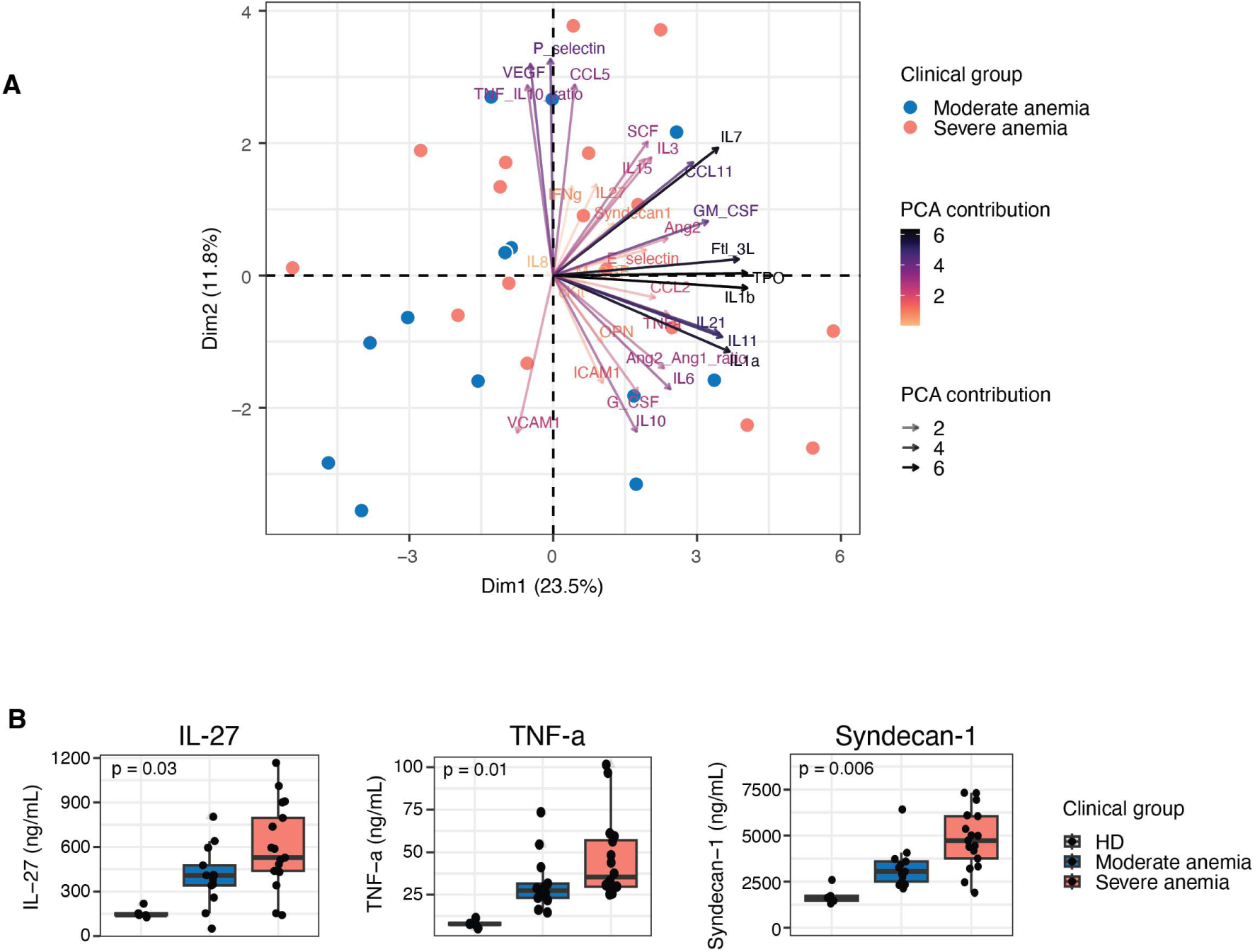
Luminex analysis of 36 protein markers in PB between SMA and MMA. **A.** Principal component analysis of Luminex markers from BM samples. **B.** Individual markers with significant difference (*p*<0.05) in PB samples between SMA and MMA.

**Figure S2.**
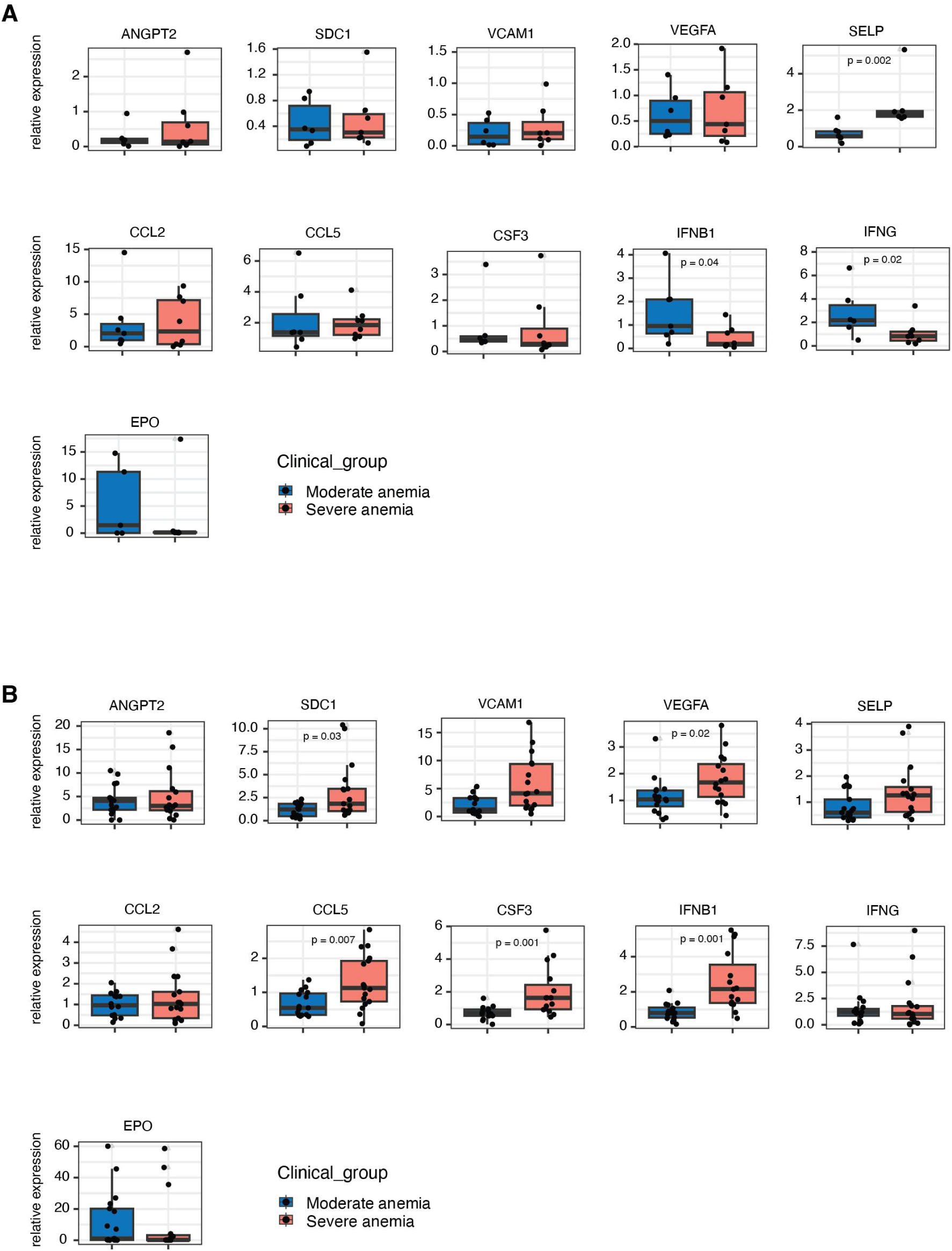
qRT-PCR analysis of 11 marker genes in PBMCs (A) and BBMCs (B). Markers overlap with their respective proteins in the Luminex panel.

**Figure S3.**
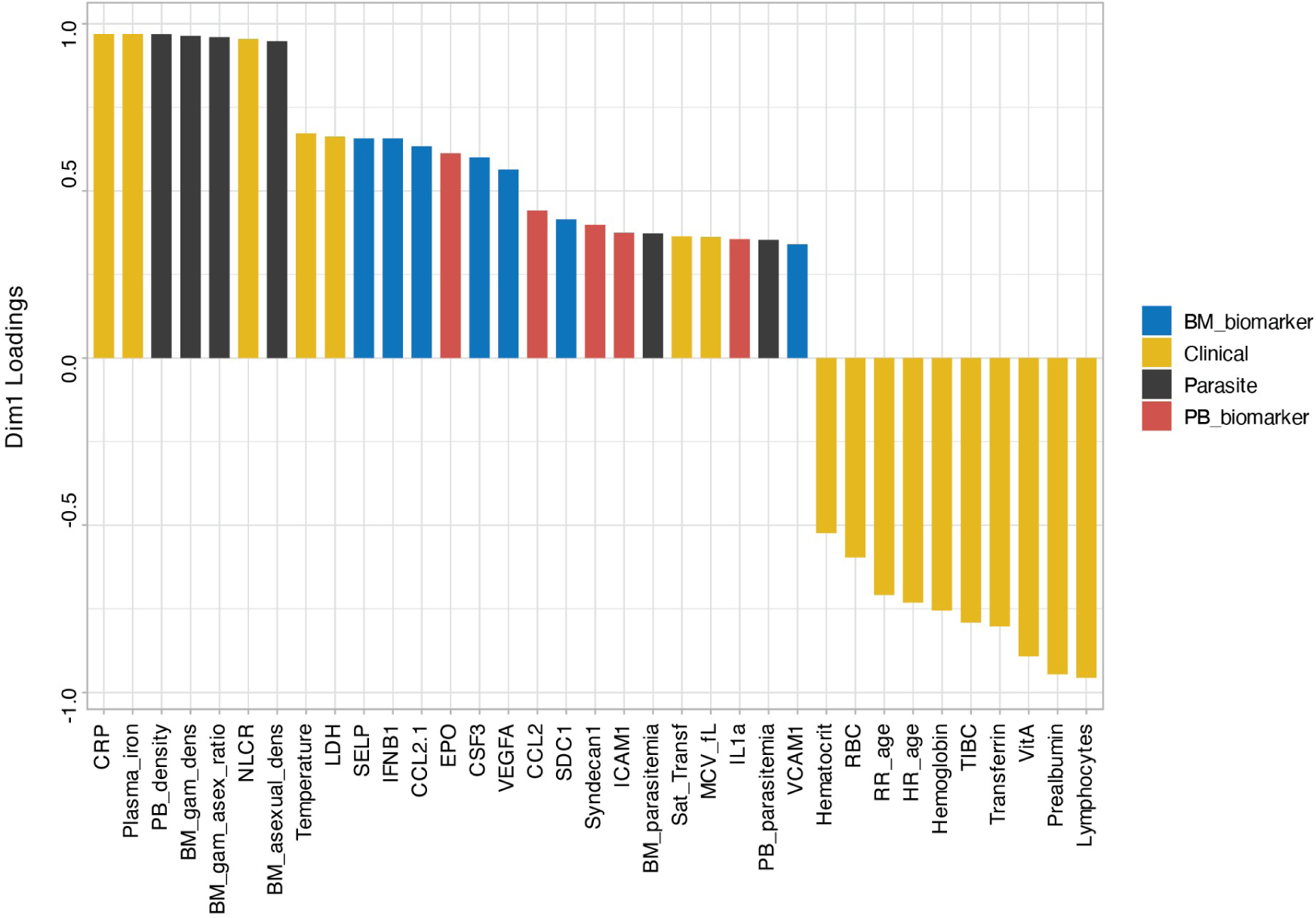
Bar blot from Multiple Factor Analysis (MFA) to integrate the clinical data, PB and BM profiling.

## Supplementary tables

**Table S1.** Patient and sample information including clinical parameters.

**Table S2.** Protein and transcript markers analysed in this study.

**Table S3.** GO function enrichment analysis based on nCounter data from BMMCs.

**Table S4.** Go function enrichment analysis based on qRT-PCR data from BMMCs.

**Table S5.** Parasite differential gene expression in SMA *vs* MMA and PB *vs* BM.

